# Male odor preference in female mice is modulated across reproductive stages via the posteroventral medial amygdala

**DOI:** 10.64898/2026.03.29.712537

**Authors:** Satomi Komada, Katsura Kagawa, Ayaka Takimoto-Inose, Soichiro Yamaguchi, Saori Yano-Nashimoto

## Abstract

Male odor induces various behavioral and physiological responses across the reproductive cycle in female mice. Although male odor preference in females is reduced during pregnancy, how it changes across later stages of the reproductive cycle, including nursing and weaning, remains unclear. Here, we found that male odor preference is lost during pregnancy and nursing. To identify the olfactory systems involved in these changes, we examined neural activity using c-Fos immunohistochemistry. Male odor exposure during nursing increased neural activity in the accessory olfactory bulb and the posteroventral medial amygdala (MeApv), a key node of the accessory olfactory system, as well as in subdivisions of the central amygdala, but not in the ventromedial hypothalamus or the bed nucleus of the stria terminalis. Finally, lesions of the MeApv prevented the loss of male preference during nursing, indicating that the MeApv is required for suppression of male preference during this stage.

## Introduction

Olfaction plays a major role in intraspecific communication including sexual, parental, and aggressive behaviors across mammals ^1–4^. In females, behavioral and physiological responses to male odors change across the reproductive cycle, including sexual maturation, the estrus cycle, pregnancy, parturition, and nursing ^4–9^. For example, in mice, exposure to male odors induces estrus in sexually mature females ^5^, whereas exposure to unfamiliar male odors during early pregnancy can lead to pregnancy termination (Bruce effect) ^7^. In addition, lactating female mice exhibit maternal aggression in response to odors from intruder males ^8,9^.

Similarly, male odor preference in female mice is also thought to vary across the reproductive cycle. Non-pregnant females typically exhibit strong attraction to male conspecifics ^10^, whereas interest in male urine is reduced during pregnancy ^11,12^. However, how male odor preference changes across later stages of the reproductive cycle, including nursing and weaning, remains unclear.

In mice, olfactory cues were mainly detected by two anatomically and functionally distinct systems: the main olfactory system and the accessory olfactory system. The main olfactory system detects volatile environmental odorants via the olfactory epithelium, with sensory signals projecting to the main olfactory bulb (MOB) ^13^. In contrast, the accessory olfactory system plays a major role in intraspecific communication by detecting pheromones and other social chemosignals. Signals generated in the vomeronasal organ are conveyed to the accessory olfactory bulb ^1,13^. Neurons in the accessory olfactory bulb (AOB) project predominantly to the medial amygdala (MeA). The MeA, in turn, sends outputs to multiple downstream regions implicated in social, reproductive, and defensive behaviors, including the bed nucleus of the stria terminalis (BNST), the ventromedial hypothalamus (VMH), and the central amygdala (CeA) ^14–19^.

These olfactory systems are regulated by estrus cycles and reproductive stages ^6,11,20^. For example, gonadectomy and replenishment of sex steroids modulate neural processing of pheromones ^20^. In addition, responsiveness of the olfactory systems to male odors changes across the estrus cycle and during early pregnancy ^6,11^. Therefore, such modulation of olfactory processing is likely to alter behavioral responses to social odors, including male-derived chemosignals. However, how such modulation affects odor preference during later reproductive stages, including lactation, remains poorly understood.

In this study, we aimed to characterize how male odor preference in female mice changes during the non-pregnant, pregnant, nursing, and post-weaning stages. To identify the olfactory systems involved in these changes, we examined neural activity in the main and accessory olfactory bulbs as well as in the downstream brain regions. We then focused on the posteroventral medial amygdala (MeApv), which was activated by male exposure during nursing and is known as a secondary center of the accessory olfactory system, and tested its functional role in male odor preference by lesioning this region.

## Materials and methods

### Animals

All experimental procedures in this study were approved by the Animal Care and Use Committee of Hokkaido University (approved protocol numbers: 20-0066 and 23-0026). All experiments were conducted in the Experimental Animal Facility of the Faculty of Veterinary Medicine, which obtained full accreditation from AAALAC International. Adult female C57BL/6JJcl mice were purchased from CLEA Japan (Tokyo, Japan) or bred in our animal facility. Mice were group-housed in standard cages (170 × 280 × 120 mm), with a maximum of seven mice per cage until the start of the experiments. The mice were kept at 22 ± 4 □with a 12:12 h light/dark cycle with light on at 7:00 with access to food and water ad libitum. All experiments were conducted during the light phase. The estrus cycle was categorized via the inspection of vaginal smears.

### Cohabitation with a male, confirmation of pregnancy and delivery, and weaning

Proestrus females were housed with a male. On the next morning, the presence of a vaginal plug was checked, after which the male was removed. The day on which a plug was detected was designated as gestational day 0 (G0). Behavioral experiments were conducted on G1-2, G6-8, and G12-15; data from G1-2 and G6-8 are presented in Supplementary Fig. 1.

To examine the effect of cohabitation with a male (Supplementary Fig. 1), diestrus females were housed with a male overnight, after which the male was removed. Estrus cycles were subsequently monitored to confirm the normal cyclicity following cohabitation.

Nursing, weaning, and non-nursing females were prepared as follows. Females were housed with a male until pregnancy was confirmed based on an increase in body weight (> 26 g) and visible abdominal distention, after which the male was removed. The day of delivery was defined as postpartum day 0 (PPD0). Experiments in nursing females were conducted on PPD6-9. For weaning females, pups were weaned on PPD21-23, and estrus cycles were monitored thereafter. For non-nursing females, pups were removed on PPD0-1.

All experiments, except those conducted in pregnant and nursing females, were performed during diestrus.

### Lesion of MeApv

Lesions of MeApv were conducted by the injection of the excitotoxic amino acid, N-methyl-D-aspartic acid (NMDA; 20 mg/ml, Sigma-Aldrich, USA) ^21^. The lesions for the nursing and weaning groups were conducted during pregnancy. Briefly, mice were anesthetized with 0.75 mg/kg medetomidine, 4.0 mg/kg midazolam, and 5.0 mg/kg butorphanol (i.p.) and fixed on stereotaxic instruments (SR-5M-HT, SM-15R, Narishige, Japan). The stereotaxic coordinate was AP −1.65 mm, ML ± 2.2mm, and VD +5.45 mm from the bregma. NMDA was injected at a volume of 40 nl per side through a glass pipette (diameter: approximately 50 μm). In sham-operated subjects, a glass pipette was inserted into the target, but NMDA was not injected. Following the surgery, the mice were awakened with atipamezole (0.75 mg/kg i.p.), and treated with enrofloxacin (10 mg/kg s.c.) and meloxicam (5 mg/kg s.c.) for 3 days. At least a week of recovery was allowed before the behavioral experiments. The lesion areas were confirmed with NeuN-stained brain sections after the experiment.

### Collection of urine

Urine was collected from group-housed C57BL/6 male and female mice older than 10 weeks by gentle abdominal massage. Urine samples were pooled from at least four animals of the same sex. The pooled urine was stored at −30 □ and used for experiments within one month of collection.

### Odor preference test

To evaluate the odor preference, odor preference tests were conducted in a home cage. The subject females were transferred to a new home cage containing an acrylic board (50 × 50 mm) and singly housed for at least one day before testing. On the test day, the acrylic board and environmental enrichment were removed, after which behavioral experiments started. For nursing females, dams with pups were transferred to a new home cage in the same manner, and pups were removed 1 hour before the experiment. Each subject was tested only once. All experiments were conducted between 7:00 and 13:00.

The experimenter applied 20 μl of odor stimuli onto filter paper (10 × 10 mm) placed on an acrylic board and then placed the board in the home cage. Odor stimuli used for analysis consisted of male and female urine. Prior to urine exposure, animals were also exposed to several non-social odorants (distilled water, mineral oil, eugenol, acetaldehyde, and ammonia); these data were not included in the present study. Behavior was video recorded for 3 minutes after each odor exposure. After the experiments, vaginal smears were collected to confirm that the estrus cycle was diestrus.

Behavior was manually scored using Solomon Coder (version 19.08.02). Sniffing was defined when the mouse directly contacted the filter paper with its nose. Periods during which the filter paper was not directly accessible to the mouse were defined as inaccessible time. These periods included instances in which the filter paper was covered by bedding or the acrylic board. The sniff rate was calculated as the proportion of sniffing time relative to the total test duration. When inaccessible periods occurred, the sniff rate was calculated by excluding the inaccessible time from the total test duration.

### Preparation of brain sections

For the c-Fos immunostaining, nursing and weaning females were used. For 2 days prior to the experiment, weaning females were transported to a new home cage and singly housed. For nursing females, dams and pups were transferred to a new cage in the same manner. For odor stimulation, the experimenter gently restrained the mice by hand and applied 30 μl of male or female urine directly to the nose and returned the animals to their home cages.

Perfusion was conducted 1.5-2 hours after the urine exposure. Under deep anesthesia, females were transcardially perfused with phosphate-buffered saline (PBS), followed by 4% (w/v) paraformaldehyde in PBS. The brains were collected and postfixed overnight with 4% paraformaldehyde at 4□ and then cryoprotected by immersing in 20% and 30% (w/v) sucrose in PBS until they sank. Then, the brains were frozen in OCT compound (Sakura finetek, Japan) and stored at −80□ until sectioning.

The brains were sectioned coronally at 40 µm using a cryostat and stored in antifreeze solution (containing 10% of 0.2 M PB, 30% glycerol, and 30% ethylene glycol (v/v)) at −20□. Every fifth section from the serial sections was processed for immunohistochemistry.

### Immunohistochemistry

Immunohistochemistry was performed largely according to a previously described protocol ^22^, with minor modifications. Free-floating brain sections were washed with PBS containing 0.1% TritonX-100 (PBST) and immersed in methanol containing 0.3% H_2_O_2_ for 5 min. The sections were then washed in PBST and blocked with 0.8% Block Ace (Megmilk Snow Bland, Japan) in PBST for 30 min at room temperature. The sections were incubated overnight at 4LJ with a rabbit monoclonal antibody against c-Fos (1:2000, 9F6, Cell Signaling Technology, USA) or a mouse monoclonal antibody anti NeuN (1:6000, MAB377, Chemicon, USA). After washing in PBST, the sections were incubated for 1 hour with a biotin-conjugated goat anti-rabbit secondary antibody (1:2000, BA-1000, Vector Laboratories, USA) or a biotin-conjugated horse anti-mouse secondary antibody (1:2000, BA-2000, Vector Laboratories). After washing in PBST, signal amplification was performed using ABC peroxidase reagent (PK-6100, Vector Laboratories) for 1 hour, washed with PBST, and then visualized with DAB solution (SK-4100, DAB Substrate Kit, Vector Laboratories) for 10 min according to the manufacturer’s instructions. The stained sections were washed with PBS and then mounted on MAS-coated slides (Matsunami, Japan). The sections stained with c-Fos were Nissl-stained. Sections were dehydrated with a series of ethanol, cleared with xylene, and mounted using a mounting medium (Softmount, Fujifilm Wako, Japan).

Photomicrographs were acquired using a virtual slide scanner (Nano Zoomer 2.0-RS, Hamamatsu Photonics, Japan). Image processing and quantification were performed using Fiji ImageJ (version 1.54f) ^23^. c-Fos-immunoreactivity was separated from Nissl staining using the Colour Deconvolution 2 plugin ^24^. The extracted c-Fos channel was binarized using a fixed intensity threshold of 120. c-Fos-positive cells were automatically quantified using the Analyze Particles function in Fiji ImageJ. The number of c-Fos-positive cells was calculated per unit area and used for statistical analyses.

### Statistical analysis

Statistical analyses were conducted using R (version 4.5.1) ^25^. Paired comparisons between two conditions were performed using the Wilcoxon signed-rank test. Unpaired comparisons between two groups were conducted by the Wilcoxon rank-sum test. Full statistical details are provided in the Supplementary Table.

## Results

### Male odor preference in female mice was lost during pregnancy and nursing

To examine whether male odor preference in female mice varies across reproductive stages, we conducted odor preference tests in non-pregnant, pregnant, nursing, and weaning females (Fig. 1A). Female mice were sequentially presented with female and male odors, and preference was quantified as the difference in sniffing time between the two odors (Fig. 1B-C). Non-pregnant females exhibited a clear preference for male over female odor, whereas this preference was not observed in pregnant females.

**Fig. 1.**
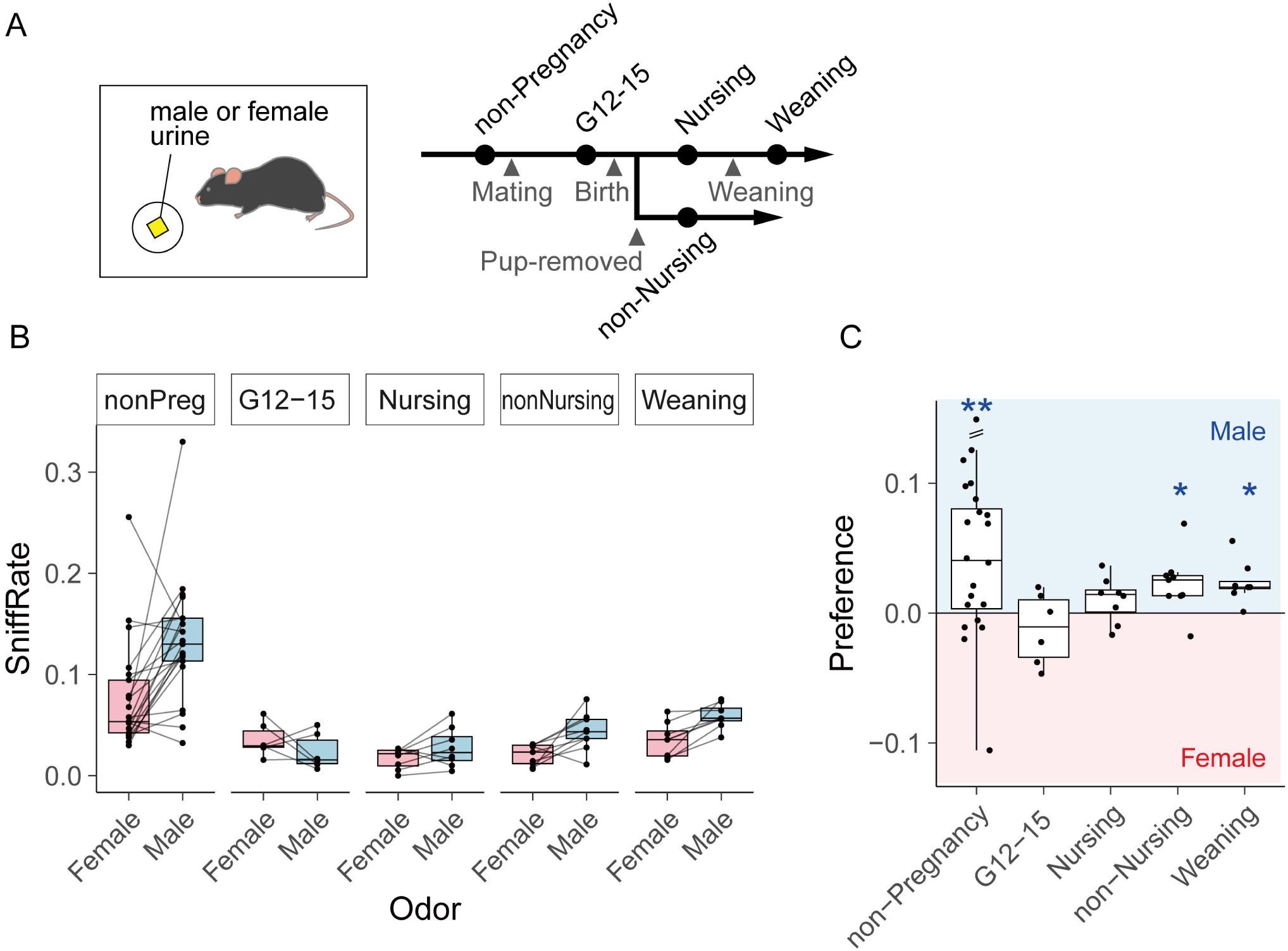
Pregnant and nursing females did not show male preference. A Schematic of the odor preference test and experimental timeline across reproductive stages. B Boxplots of the sniff rate (total sniffing duration divided by test duration). Each dot represents an individual trial, and dots from the same mouse are connected by lines. C Boxplots of the preference (the male sniffing rate minus the female sniffing rate). Non-pregnant females showed a significant preference for male urine. This male preference was absent during pregnancy and nursing, but re-emerged after pup removal (non-nursing and weaning). Asterisks indicate that the preference is significantly different from zero (Wilcoxon signed-rank test; \**p* < 0.05, \*\**p* < 0.01). See also Supplemental Fig. 1.

Male odor preference was transiently reduced immediately after cohabitation with a male, regardless of pregnancy status (Supplementary Fig. 1). Therefore, changes in odor preference during early pregnancy could not be clearly dissociated from the effects of recent cohabitation. However, this cohabitation-induced reduction in male preference was no longer observed 12-15 days after cohabitation. Accordingly, the loss of male preference observed during mid-pregnancy (G12-15) reflects pregnancy-related changes rather than residual effects of male cohabitation.

Following pregnancy, male odor preference was also absent during nursing. In contrast, male preference recovered after weaning (Fig. 1C). A similar recovery was observed in postpartum females whose pups were removed just after birth. These results indicate that the recovery of male preference is associated with the termination of nursing, rather than simply with the passage of time after male cohabitation and parturition.

### Male urine exposure during nursing induced activation of the accessory olfactory system

To examine whether stage-dependent variation in male odor preference is associated with neural activity in the main or accessory olfactory pathways, we assessed odor-evoked neural activation in the main olfactory bulb (MOB) and the accessory olfactory bulb (AOB) in nursing and weaning females (Fig. 2A-B). The main olfactory system processes general volatile odors via the MOB, whereas the accessory olfactory system detects pheromonal cues through the vomeronasal organ and AOB. Neuronal activity was evaluated by c-Fos immunohistochemistry following exposure to male urine, with female urine used as a control odor (Fig. 2C-I).

**Fig. 2.**
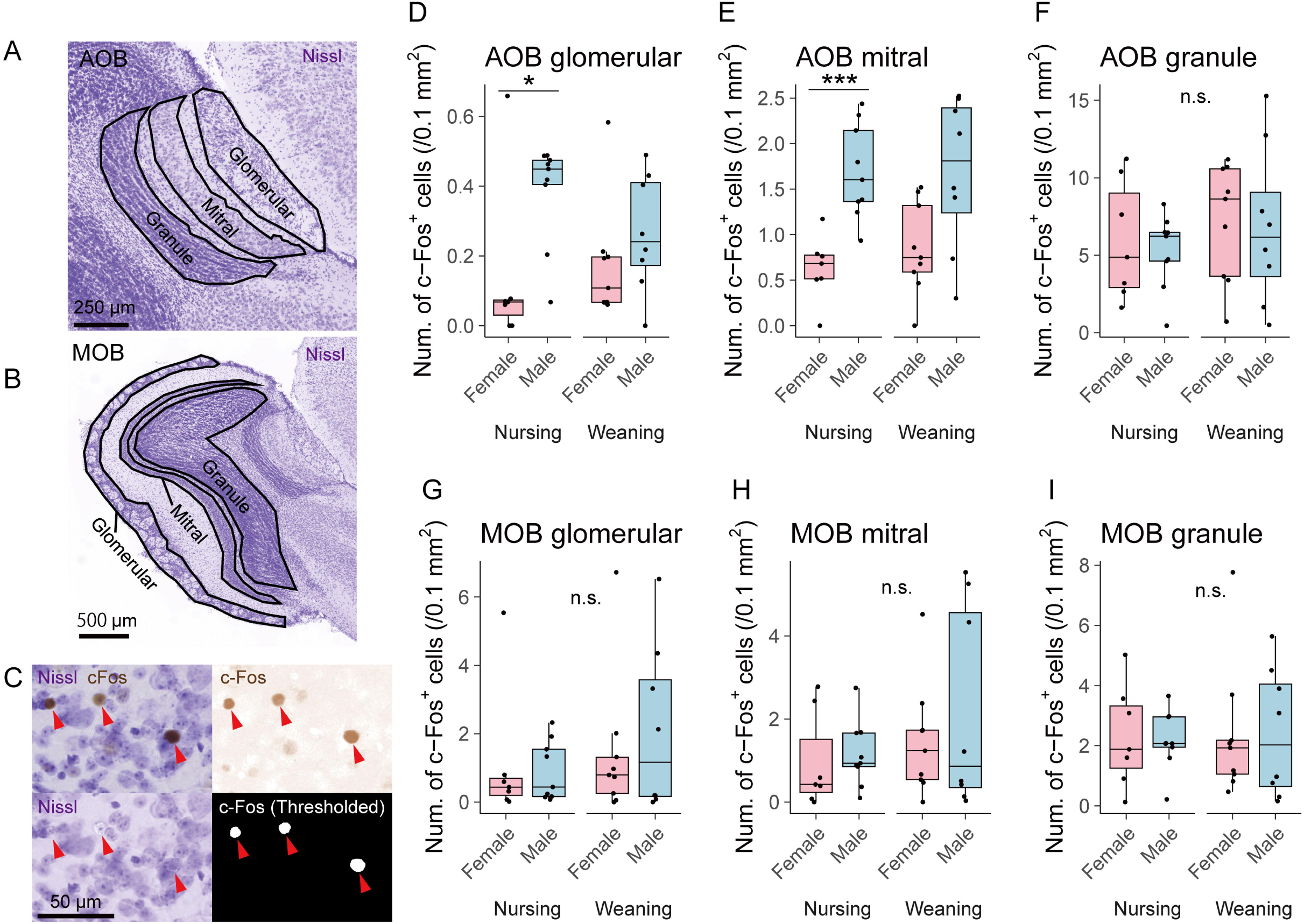
Male urine exposure activated the AOB but not the MOB in nursing females. A-B Representative Nissl-stained sections of the AOB (A) and the MOB (B) with anatomical layers indicated. C Representative images showing c-Fos immunoreactivity, the c-Fos channel extracted by color deconvolution, and thresholded c-Fos signals used for quantification. D-I Boxplots of the number of c-Fos-immunoreactive cells in the AOB (D-F) and the MOB (G-I) after exposure to male and female urine. In the glomerular and mitral cell layers of the AOB in nursing females, the number of c-Fos immunoreactive cells increased after male urine exposure. Asterisks indicate significant differences between female and male urine exposure (Wilcoxon rank-sum test; \**p* < 0.05, \*\*\**p* < 0.001).

Male urine exposure increased c-Fos expression in the AOB of nursing females, whereas no such increase was observed in weaning females (Fig. 2D-F). This increase in c-Fos expression was particularly prominent in the glomerular and mitral layer but not in the granule layer of AOB. In contrast, male urine exposure did not induce significant changes in c-Fos expression in MOB in either nursing or weaning females (Fig. 2G-I).

Given that male urine selectively activated AOB during nursing, we next examined the downstream regions of the accessory olfactory system (Fig. 3, Supplementary Fig. 2). Neurons in the mitral cell layer of AOB project to the medial amygdala (MeA). Among MeA subregions, the posteroventral subregion (MeApv) was selectively activated by male urine exposure during nursing but not in weaning females (Fig. 3C). In contrast, other subregions of MeA did not show significant changes in neuronal activity in nursing females (Fig. 3D, Supplementary Fig. 2A-B).

**Fig. 3.**
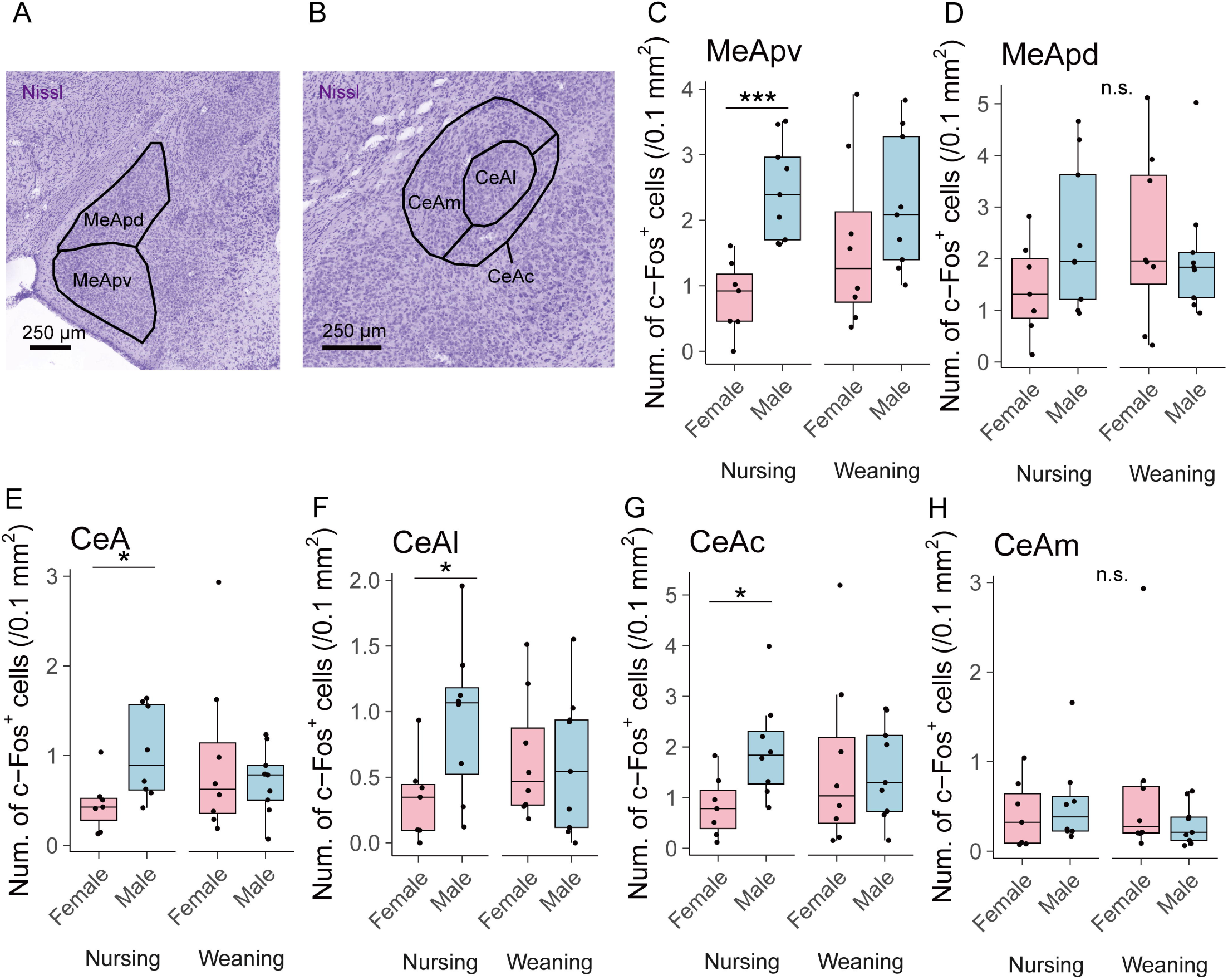
Male urine exposure activated the MeApv, CeAl, and CeAc in nursing females. A-B Representative Nissl-stained sections of the MeA (A) and the CeA (B) with anatomical subregions indicated. C-H Boxplots of the number of c-Fos-immunoreactive cells in the MeA (C-D) and the CeA (E-H) after exposure to male or female urine. In the MeApv of nursing females, male urine exposure increased the number of c-Fos immunoreactive cells, whereas no comparable changes were observed in other MeA subregions or in post-weaning females. Similarly, in the CeA of nursing females, male urine exposure increased the number of c-Fos-immunoreactive cells. Such changes were observed only in the CeAl and CeAc. Asterisks indicate significant differences between female and male urine exposure (Wilcoxon rank-sum test; \**p* < 0.05, \*\*\**p* < 0.001). See also Supplemental Fig. 2.

Given the selective activation of MeApv during nursing, we next examined downstream regions that receive projections from MeA, including the ventromedial hypothalamus (VMH), the bed nucleus of the stria terminalis (BNST), and the central amygdala (CeA). Male urine exposure during nursing significantly increased neuronal activity in CeA (Fig. 3E). Within CeA, increased c-Fos expression was observed in the lateral and capsular subdivisions (CeAl and CeAc), but not in the medial subdivision (CeAm), indicating that CeA activation during nursing was accounted for by increased activity in CeAl and CeAc (Fig. 3F-H). In contrast, male urine exposure during nursing did not induce significant changes in neuronal activity in BNST or VMH (Supplementary Fig. 2C-F).

### The loss of male preference during nursing required MeApv

To determine whether MeApv is required for the loss of male odor preference during nursing, we examined the effect of N-methyl-D-aspartate (NMDA) lesions in MeApv on odor preference behavior (Fig. 4, Supplementary Fig. 3). In sham-operated nursing females, male odor preference was absent, consistent with the results in the intact females (Fig. 1B-C, 4B). In contrast, nursing females with MeApv lesions retained a significant preference for male over female odor (Fig. 4B), indicating that disruption of MeApv prevented the loss of male preference during nursing.

**Fig. 4.**
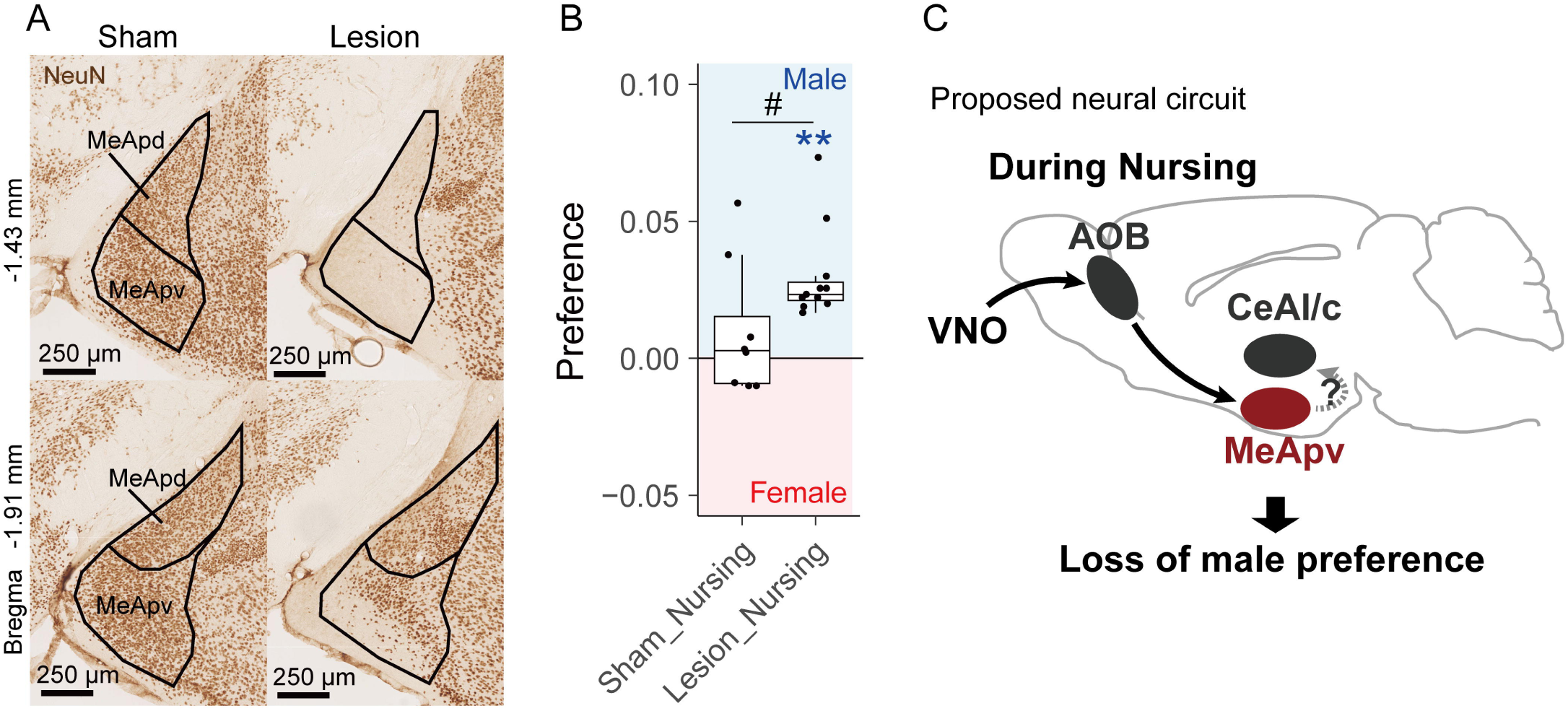
NMDA lesion in the MeApv inhibited male preference in nursing females. A Representative lesion area following NMDA injection into the MeApv. Sections were stained with the mature neuronal marker NeuN. Lesion animals included cases with partial damage extending beyond the MeApv and/or unilateral lesions. B Boxplots of the preference (the male sniffing rate minus the female sniffing rate). Whereas the nursing females with Sham operation did not show male preference, the lesioned females showed male preference even during nursing. The preference was significantly different between the lesioned and sham-operated animals. Asterisks indicate that the preference is significantly different from zero (Wilcoxon signed-rank test; \*\**p* < 0.01). The hash symbol (#) indicates the significant difference between the lesion and sham groups (Wilcoxon rank-sum test; #*p* < 0.05). See also Supplemental Fig. 3.

## Discussion

In this study, we demonstrated that male odor preference in female mice varies across reproductive stages and is lost during pregnancy and nursing. We further show that male urine exposure during nursing selectively activates the accessory olfactory system, as evidenced by increased neuronal activity in the AOB, MeApv, CeAl, and CeAc. Importantly, lesions of MeApv prevented the loss of male preference during nursing, indicating that MeApv is required for the suppression of male preference during nursing.

Although the loss of male preference was consistently observed during pregnancy and nursing, the regulatory mechanisms underlying this behavioral change may differ across reproductive stages. In the present study, we focused specifically on the nursing period and examined neural activity and causal involvement of MeApv using c-Fos immunohistochemistry and lesion experiments. By contrast, the mechanisms responsible for the loss of male preference during pregnancy were not directly addressed here. Given differences in hormonal status and infant-derived sensory stimulation between pregnancy and nursing, distinct neural and endocrine processes may regulate male odor preference at these stages. Further studies will be required to determine whether the neural mechanisms underlying the loss of male preference during nursing are shared with, or distinct from, those operating during pregnancy.

Our findings support the involvement of an accessory olfactory circuit in the loss of male odor preference during nursing. The accessory olfactory system is generally thought to process social and pheromonal cues, and the present results are consistent with this view ^1,13^. Notably, male urine exposure during nursing consistently activated the AOB, MeApv, CeAl, and CeAc. Within the accessory olfactory system, sensory signals detected by the vomeronasal organ are transmitted to the AOB and subsequently to the MeA, which in turn projects to multiple downstream brain regions involved in social behavior ^1^. Anatomical studies have shown that neurons in the MeA project to the CeA, including the CeAl and CeAc; however, projections from the MeApv are considered relatively sparse (Pardo-Bellver et al., 2012). Accordingly, although direct projections from the MeApv to the CeAl and CeAc appear to be weak, functional interactions between these regions may contribute, at least in part, to the loss of male odor preference during nursing. Such interactions could be mediated by indirect pathways or polysynaptic circuits, and further studies will be required to clarify the precise anatomical and functional connectivity underlying this effect.

CeA has been widely implicated in the processing of aversive, defensive, and negatively valenced information, and its activation is often associated with avoidance-related behaviors ^26^. Therefore, it is possible that the activation of CeA during nursing enhances avoidance-related processing of male odor and thereby suppresses male odor preference typically observed in non-pregnant females. Notably, other downstream targets of MeA, such as BNST and VMH, which are also commonly linked to avoidance, defensive, or aggressive behaviors, were not significantly activated by male urine exposure during nursing ^17,27,28^. This selective activation of CeA, but not BNST or VMH, suggests that suppression of male preference during nursing may rely on a specific accessory olfactory pathway rather than on a broad activation of defensive or threat-related circuits.

Avoidance of male mice during nursing is likely advantageous for female reproductive success. With the exception of a brief postpartum estrus shortly after parturition, lactating females do not ovulate and thus have limited opportunities for mating and subsequent pregnancy ^29^. In contrast, encounters with unfamiliar males increase the risk of infanticide, as sexually inexperienced males are more likely to attack infants ^30^. Thus, male avoidance during nursing may function as an adaptive strategy to enhance offspring survival. Our results further suggest that neural circuits in the accessory olfactory pathway, including the MeApv, may contribute to reproductive-state-dependent modulation of social responses to male cues. Such state-dependent plasticity in olfactory circuits may represent a general mechanism by which mammalian females adapt their social behaviors to changing reproductive demands.

## Supplementary figure legends

**Supplementary Fig. 1.**
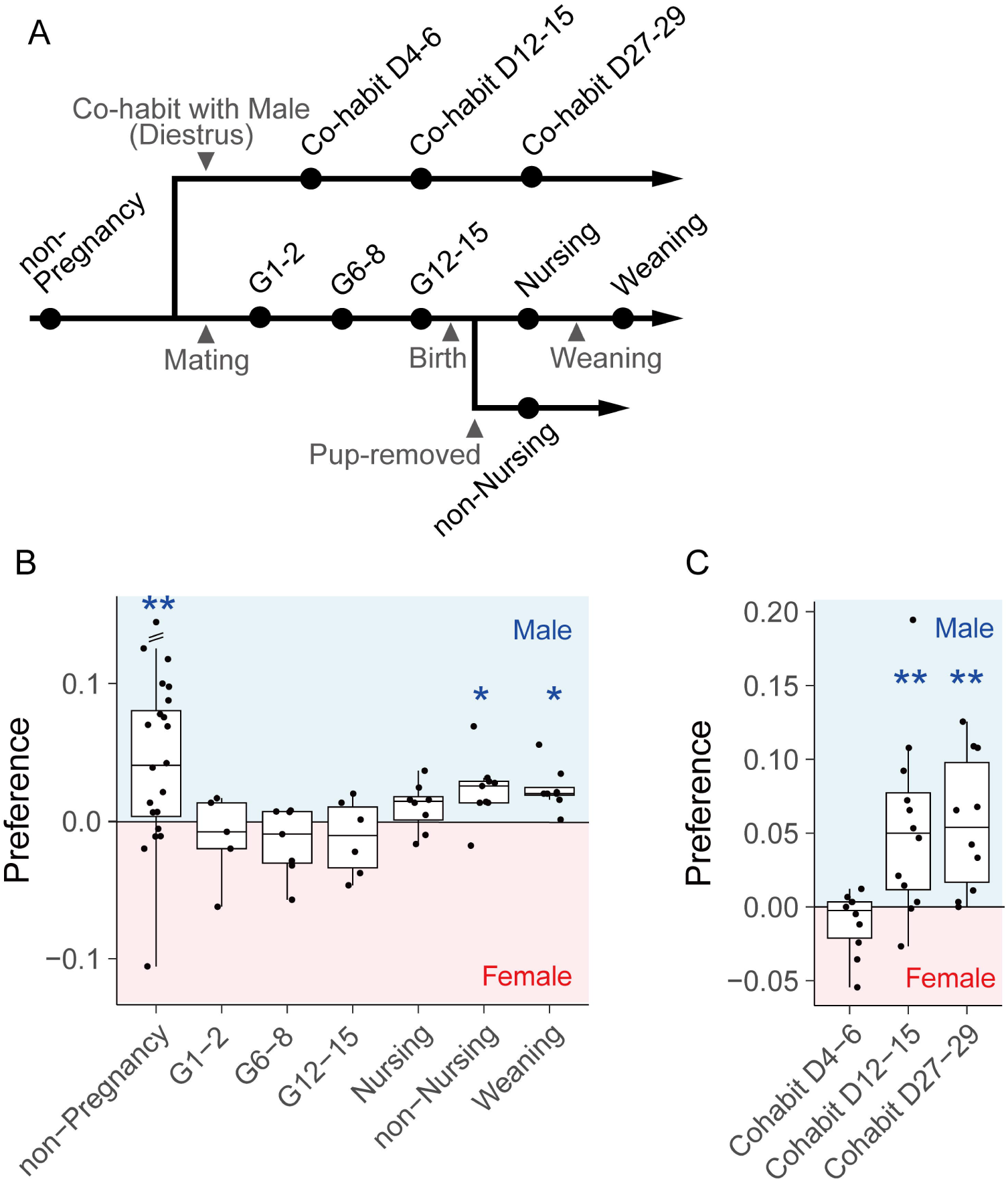
Odor preference in female mice across reproductive and experimental conditions. Data includes all conditions shown in Fig. 1, as well as additional conditions not displayed in the main figure. A Schematic of the odor preference test and experimental timeline across reproductive stages. B Boxplots of the preference (the male sniffing rate minus the female sniffing rate). C Boxplots of the preference 4-6, 12-15, or 27-29 days after cohabitation with a male mouse during (Cohabit D4-6, D12-15, D27-29). Asterisks indicate that the preference is significantly different from zero (Wilcoxon signed-rank test; \**p* < 0.05, \*\**p* < 0.01).

**Supplementary Fig. 2.**
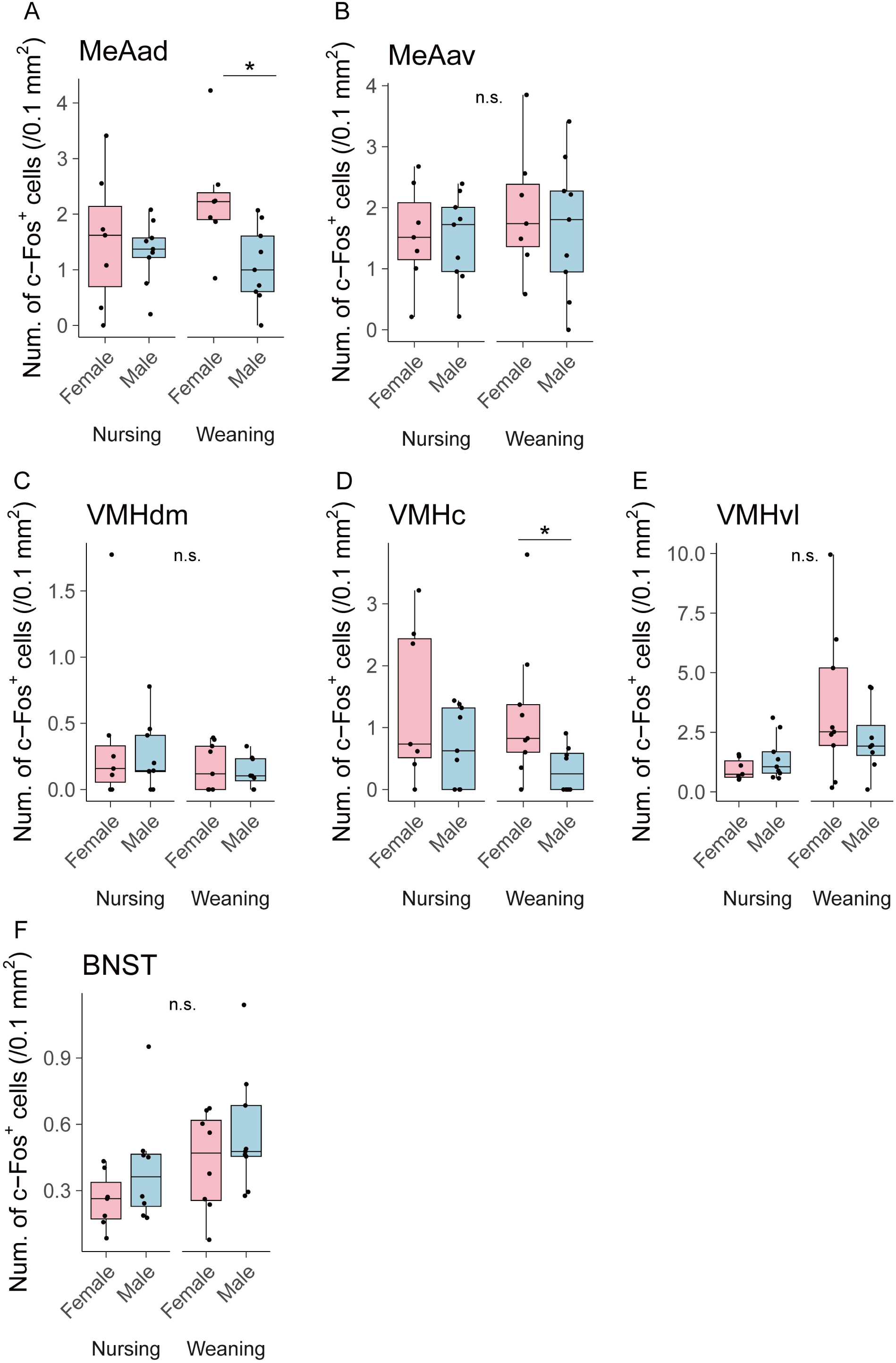
Male urine exposure did not activate the MeAa, VMH, and BNST in nursing females. A-F Boxplots of the number of c-Fos-immunoreactive cells in the MeAa (A-B), the VMH (C-E), and BNST (F) after exposure to male or female urine. In these brain regions, no significant increase in c-Fos immunoreactive cells was observed. Asterisks indicate significant differences between female and male urine exposure (Wilcoxon rank-sum test; \**p* < 0.05). Data are from the same experiments as shown in Fig. 3.

**Supplementary Fig. 3.**
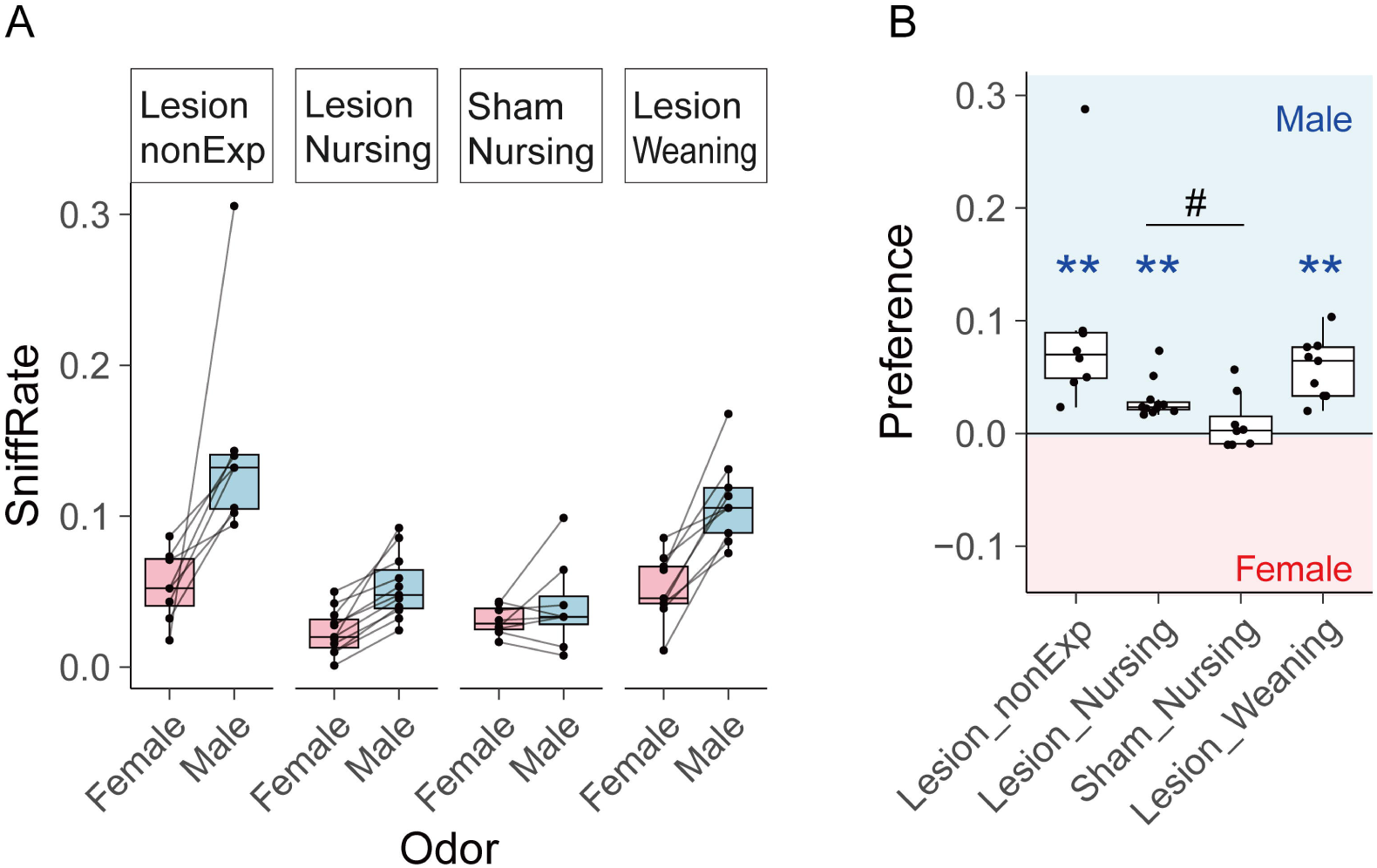
Odor preference across reproductive stages in females with the MeApv lesions. Data includes all conditions shown in Fig. 4, as well as additional conditions not displayed in the main figure. A Boxplots of the sniff rate (total sniffing duration divided by test duration). Each dot represents an individual trial, and dots from the same mouse are connected by lines. B Boxplots of the preference (the male sniffing rate minus the female sniffing rate). Asterisks indicate that the preference is significantly different from zero (Wilcoxon signed-rank test; \*\**p* < 0.01). The hash symbol (#) indicates the significant difference between the lesion and sham groups (Wilcoxon rank-sum test; #*p* < 0.05).

## Author contributions

S.Y.-N. and S.K. conceived and organized the study.

S.K. and S.Y.-N. designed and performed the experiments, with assistance from K.K.

S.Y. provided support for the study organization.

S.Y.-N. and A.T.-I. contributed to funding acquisition.

S.Y.-N. wrote the manuscript with contributions from all the authors.

All authors discussed the results and approved the final version of the manuscript.

## Acknowledgments

We thank all staff managing the experimental animal facility in the Faculty of Veterinary Medicine at Hokkaido University and the Open Facility of the Global Facility Center at Hokkaido University.

## Declaration of generative AI and AI-assisted technologies in the writing process

During the preparation of this work, the authors used ChatGPT and Grammarly in order to improve language and readability. After using these services, the authors reviewed and edited the content as needed and take full responsibility for the content of the publication.

## References

1. Murata, K., Itakura, T. & Touhara, K. Neural basis for pheromone signal transduction in mice. Front. Neural Circuits 18, 1409994 (2024).

2. Chamero, P., Leinders-Zufall, T. & Zufall, F. From genes to social communication: molecular sensing by the vomeronasal organ. Trends Neurosci. 35, 597–606 (2012).

3. Raam, T. & Hong, W. Organization of neural circuits underlying social behavior: A consideration of the medial amygdala. Curr. Opin. Neurobiol. 68, 124–136 (2021).

4. Liberles, S. D. Mammalian pheromones. Annu. Rev. Physiol. 76, 151–175 (2014).

5. Jemiolo, B., Harvey, S. & Novotny, M. Promotion of the Whitten effect in female mice by synthetic analogs of male urinary constituents. Proc. Natl. Acad. Sci. U. S. A. 83, 4576–4579 (1986).

6. Yano, S., Sakamoto, K. Q. & Habara, Y. Female mice avoid male odor from the same strain via the vomeronasal system in an estrogen-dependent manner. Chem. Senses 40, 641–648 (2015).

7. Bruce, H. M. An exteroceptive block to pregnancy in the mouse. Nature 184, 105 (1959).

8. Lonstein, J. S. & Gammie, S. C. Sensory, hormonal, and neural control of maternal aggression in laboratory rodents. Neurosci. Biobehav. Rev. 26, 869–888 (2002).

9. Martín-Sánchez, A. et al. From sexual attraction to maternal aggression: when pheromones change their behavioural significance. Horm. Behav. 68, 65–76 (2015).

10. Asaba, A., Hattori, T., Mogi, K. & Kikusui, T. Sexual attractiveness of male chemicals and vocalizations in mice. Front. Neurosci. 8, 231 (2014).

11. Serguera, C., Triaca, V., Kelly-Barrett, J., Banchaabouchi, M. A. & Minichiello, L. Increased dopamine after mating impairs olfaction and prevents odor interference with pregnancy. Nat. Neurosci. 11, 949–956 (2008).

12. Miller, C. H. et al. Pregnancy modulates responses to male odors in house mice. Horm. Behav. 175, 105802 (2025).

13. Munger, S. D., Leinders-Zufall, T. & Zufall, F. Subsystem organization of the mammalian sense of smell. Annu. Rev. Physiol. 71, 115–140 (2009).

14. Abellán-Álvaro, M., Martínez-García, F., Lanuza, E. & Agustín-Pavón, C. Inhibition of the medial amygdala disrupts escalated aggression in lactating female mice after repeated exposure to male intruders. Commun. Biol. 5, 980 (2022).

15. Liu, M., Kim, D.-W., Zeng, H. & Anderson, D. J. Make war not love: The neural substrate underlying a state-dependent switch in female social behavior. Neuron 110, 841–856.e6 (2022).

16. Demir, E. et al. The pheromone darcin drives a circuit for innate and reinforced behaviours. Nature 578, 137–141 (2020).

17. Miller, S. M., Marcotulli, D., Shen, A. & Zweifel, L. S. Divergent medial amygdala projections regulate approach-avoidance conflict behavior. Nat. Neurosci. 22, 565–575 (2019).

18. Pardo-Bellver, C., Cádiz-Moretti, B., Novejarque, A., Martínez-García, F. & Lanuza, E. Differential efferent projections of the anterior, posteroventral, and posterodorsal subdivisions of the medial amygdala in mice. Front. Neuroanat. 6, 33 (2012).

19. Ito, M. et al. The parabrachial-to-amygdala pathway provides aversive information to induce avoidance behavior in mice. Mol. Brain 14, 94 (2021).

20. Baum, M. J. & Bakker, J. Roles of sex and gonadal steroids in mammalian pheromonal communication. Front. Neuroendocrinol. 34, 268–284 (2013).

21. Tsuneoka, Y. et al. Functional, anatomical, and neurochemical differentiation of medial preoptic area subregions in relation to maternal behavior in the mouse. J. Comp. Neurol. 521, 1633–1663 (2013).

22. Anan, M., Kagawa, K., Okamatsu-Ogura, Y., Yamaguchi, S. & Yano-Nashimoto, S. Human chorionic gonadotropin inhibits locomotion but not food intake independently of the area postrema and the nucleus tractus solitarius in female mice. Physiol. Behav. 303, 115151 (2026).

23. Schindelin, J., et al. Fiji: an open-source platform for biological-image analysis. Nat. Methods 9, 676–682 (2012).

24. Landini, G., Martinelli, G. & Piccinini, F. Colour deconvolution: stain unmixing in histological imaging. Bioinformatics (2020) doi:10.1093/bioinformatics/btaa847.

25. R Core Team. R: A language and environment for statistical computing. Foundation for Statistical Computing, Vienna, Austria (2013).

26. Moscarello, J. M. & Penzo, M. A. The central nucleus of the amygdala and the construction of defensive modes across the threat-imminence continuum. Nat. Neurosci. 25, 999–1008 (2022).

27. Wang, L., Chen, I. Z. & Lin, D. Collateral Pathways from the Ventromedial Hypothalamus Mediate Defensive Behaviors. Neuron 85, 1344–1358 (2015).

28. Nordman, J. C. et al. Potentiation of divergent medial amygdala pathways drives experience-dependent aggression escalation. J. Neurosci. 40, 4858–4880 (2020).

29. Hackwell, E. C. R., Ladyman, S. R., Brown, R. S. E. & Grattan, D. R. Mechanisms of lactation-induced infertility in female mice. Endocrinology 164, bqad049 (2023).

30. Tachikawa, K. S., Yoshihara, Y. & Kuroda, K. O. Behavioral transition from attack to parenting in male mice: a crucial role of the vomeronasal system. J. Neurosci. 33, 5120–5126 (2013).

